# The Role of Interleukin-1 cytokine family (IL-1β, IL-37) and interleukin-12 cytokine family (IL-12, IL-35) in eumycetoma infection pathogenesis

**DOI:** 10.1101/502609

**Authors:** Amir Abushouk, Amre Nasr, Emad Masuadi, Gamal Allam, Emmanuel E. Siddig, Ahmed H. Fahal

**Author notes:** **Corresponding Author** Professor Ahmed Hassan Fahal, Professor of Surgery, The Mycetoma Research Centre, University of Khartoum, Khartoum, Sudan. E. mail.

## Abstract

Mycetoma is a neglected tropical disease, endemic in many tropical and subtropical regions, characterised by massive deformity and disability and can be fatal if untreated early and appropriately. Interleukins (IL)-35 and IL-37 are newly discovered cytokines that play an important role in suppressing the immune system. However, the expression of these interleukins in patients with *Madurella mycetomatis (M. mycetomatis)* induced eumycetoma has not yet been explored. This study aims to determine the levels of the IL-1 family (IL-1β, IL-37) and IL-12 family (IL-12, IL-35) in a group of these patients and the association between these cytokines levels and the patients’ demographic characteristics. The present, a case-control study was conducted at the Mycetoma Research Centre, Soba University Hospital, University of Khartoum, Sudan and it included 140 individuals. They were divided into two groups; group I: healthy controls [n = 70; median age 25 years (range 12 to 70 years)]. Group II: mycetoma patients [n = 70 patients; median age 25 (range 13 to 70 years)]. Cytokines levels were measured in sera using enzyme-linked immunosorbent assay (ELISA).

There was no significant correlation between the IL-1β and IL-12 levels and the lesions size and disease duration, whereas levels of IL-37 and IL-35 were significantly correlated with that. The analysis of the risk factors of higher circulatory levels of IL-37 in patients of mycetoma showed a significant negative association with IL-1β cytokine, where a unit increment in IL-1β will decrease the levels of IL-37 by 35.28 pg/ml. The levels of IL-37 among the patients with a duration of mycetoma infection ≤ one year had significantly decreased by an average of 18.45 compared to patients with a mycetoma infection’s duration of ≥ 5years (reference group). Furthermore, the risk factors of higher levels of IL-35 in mycetoma patients revealed a significant negative association with IL-12, as a unit increment in IL-12 decreases the levels of IL-35by 8.99 pg/ml (*p* < 0.001). Levels of IL-35 among the patients with duration of mycetoma infection ≤ one year had significantly decreased (*p*-value = 0.002) on average by 41.82 compared to patients with a duration of mycetoma infection ≥ five years (reference group). In conclusion, this study indicates that both IL-35 and IL-37 are negatively associated with the levels of IL-1β and IL-12 in eumycetoma mycetoma infection; and high levels of IL-37 and IL-35 may have a negative impact on disease progression.

**Authors Summary:** Mycetoma is a progressive chronic granulomatous fungal or bacterial infection that may result in massive destruction of subcutaneous tissues, muscles and bones. Mycetoma is a neglected disease which is endemic in many tropical and subtropical areas. If the disease is not treated properly, eventually it ends up with amputation and adverse medical, health and socioeconomic effects on patients and the community.

Previous data suggested a crucial role of adaptive immunity in host resistance to causative agents and the disease progression. The recently identified IL-35 and IL-37 cytokines revealed an important role in immune suppression. Nevertheless, the expression of these interleukins in patients with mycetoma has not yet been investigated. Therefore, the present case-control study aimed to determine the levels of IL-1 family (IL-1β, IL-37) and IL-12 family (IL-12, IL-35) in these patients and the association between these cytokines levels and the patients’ demographic characteristics.

The results of this study showed that the levels of IL-37 and IL-35 were consistently positively correlated with different diameters of mycetoma lesions as well as its duration. However, the levels of IL-1β and IL-12 were consistently negatively correlated with different diameters of lesions and the duration of mycetoma infection. The analysis of the risk factors of higher circulatory levels of IL-37 in patients of mycetoma showed a significant negative association with IL-1β cytokine. Furthermore, the risk factors of higher levels of IL-35 in patients of mycetoma revealed a significant negative association with IL-12. These findings uncover a possible the role of IL-35 and IL-37 in the pathogenesis of mycetoma and may declare their potential value in the treatment of mycetoma.

## Introduction

Mycetoma is a chronic granulomatous subcutaneous inflammatory disease, caused by certain bacteria (actinomycetoma) or fungi (eumycetoma). This infection progresses to affect the deep structures and bones leading to massive destruction, deformities and disabilities [1]. It constitutes a major health problem in many tropical and subtropical countries, and it is highly endemic in Sudan, Mexico, and India. In Sudan, more than 8500 patients were managed at the Mycetoma Research Centre in Khartoum, of whom 70% were infected with the fungus *M. mycetomatis.* The disease affects all age groups, but it occurs most commonly in young men at the age group 20 to 40 years [2]. The disease is usually painless, and the clinical diagnosis is commonly based on the presence of subcutaneous mass, multiple sinuses and seropurulent discharge with grains. Treatment opportunities comprise of various chemotherapeutics agents and wide surgical excision of the infected tissues and may possibly end up with limb amputation [3].

Although most individuals in endemic areas have antibodies against the causative agent of mycetoma; only a few develop the disease [4]. This disparity in host response is due to the interplay between the host and the pathogen [4]. Both innate and adaptive immunity play a role in host resistance to causative agents and the development of the disease. Therefore, T-cell responses seem to be important in the progress of mycetoma [5, 6]. A Th2-like response was reported in primary lesions and in draining lymph nodes in patients with *Streptomyces somaliensis* infection and after stimulation of peripheral blood mononuclear cells by *M. mycetomatis* antigens [5, 7], while the Th1 response was reported in the acute phase of infection and healthy endemic controls [8, 9]. Macrophages stimulated with live conidia of Pseudallescheria boydii also induced a Th2 response, whereas hyphae induced a Th1 response [10]. Experimental infection of nude athymic rats and mice with *Nocardia (N.) asteroides* let to fatal disease dissemination [11, 12]. In addition, T lymphocytes from previously immunised animals directly killed *N. asteroides* [12, 13]. Moreover, Trevino-Villarreal and associates [14] reported that *N. brasiliensis* cell wall-associated lipids are implicated in the development of experimental actinomycetoma and act principally by inhibiting several microbicidal effects of macrophages, including the inhibition of TNF-α production, phagocytosis, production of nitric oxide (NO), and bacterial killing. Also they demonstrated that *the N. brasiliensis* wall-associated lipids suppressed the expression of major histocompatibility complex class II (MHC II), CD80, and CD40 by dendritic cells (DCs) and strongly induced the production of TGF-β by these cells. It has been suggested that pre-existing Th2 environment caused by schistosomiasis promotes the development of mycetoma as patients with mycetoma were significantly more positive for schistosoma antibodies than healthy endemic controls [9]. These findings suggested that Th2 like response and anti-inflammatory/immunosuppressive cytokines could have a negative impact on mycetoma development and disease progression.

IL-1 is a polypeptide which has two forms; IL-1α and IL-1β. It is involved in the acute-phase response and is accountable for several alterations that are related to the onset of various medical disorders [15, 16]. It is demonstrated recently that higher levels of IL-1β cytokine are strongly associated with surgically treated mycetoma patients, in comparison to those treated without surgery [17]. It is known that IL-1β is a pro-inflammatory cytokine that is involved in cell death coordination [18]. IL-1β cytokine is cleaved into the mature, active form primarily by inflammasome-dependent caspase activity [18]. It is possible that mature IL-1β secretion by macrophages activates IL-1R1 on macrophages, fibroblasts and epithelial cells, inducing production of the CXC chemokine CXCL1/KC, which binds to CXCR2 on neutrophils and mediates recruitment of neutrophils from peripheral blood to stimulate inflammation at the site of mycetoma invasion. Therefore, these higher levels of IL-1β cytokine advocate a crucial role in *M. mycetomatis* pathogenesis.

IL-37, which is a member of the IL-1 family, has emerged as a potent anti-inflammatory cytokine that suppresses both innate and adaptive immune responses [19]. Its role in human diseases is not completely understood yet [20]. However, the anti-inflammatory properties of IL-37 have been associated with inflammatory diseases, such as systemic lupus erythematosus (SLE) [21], and inflammatory bowel disease [22]. It has been reported that IL-37 is negatively associated with pro-inflammatory cytokines such as IL-1β, IL-6, IL-17, TNF-α and IFN-γ in peripheral blood mononuclear cells (PBMCs) of patients with degenerative intervertebral discs [23] and Graves’ disease (GD) [24]. IL-37 protein level in PBMCs and dendritic cells (DCs) is up-regulated when stimulated by Toll-like receptor (TLR) ligands or pro-inflammatory cytokines [25]. In vitro, overexpression of IL-37 in macrophages or epithelial cells greatly inhibits the production of major pro-inflammatory cytokines such as TNF-α, IL-1α, IL-1β, IL-6, IFN-γ and macrophage inflammatory protein 2 [25, 26]. In vivo, IL-37 transgene protects mice from lipopolysaccharide-induced shock and chemical-induced colitis [19, 27]. IL-37 interferes with the innate protective anti-Candida host response by reducing the production of pro-inflammatory cytokines and suppressing neutrophil recruitment in response to Candida infection, resulting in increased susceptibility to disseminated candidiasis [28]. Moreover, IL-37 markedly reduced inflammasome activation and disease severity in murine aspergillosis [29]. In addition to its role in innate immunity; IL-37 plays a pivotal role in regulating adaptive immunity by inducing regulatory T (T_reg_) cells and impairing activation of effector T-cell responses [30]. To our knowledge, there has been no study to report the relationship between IL-37 and mycetoma pathogenesis so far.

Interleukin-12 (IL-12) is frequently denoted as a B cell cytokine, although it is mainly formed by innate immune cells, comprising epithelial cells, DCs, and macrophages [31]. IL-12 is a multimer that plays a fundamental role in immune regulation and is extensively involved in infections. It binds to the heterodimeric IL-12 receptor, which is principally present in T cells and on natural killer (NK) cells [31]. IL-12 induces Th1 responses [32], which consequently increase the cytotoxic cytokines, in addition to IFN-γ by T cells [31, 32].

IL-35 is a recently identified heterodimeric cytokine which belongs to the IL-12 cytokine family, composed of the subunits of IL-27; β chain Epstein-Barr-virus (EBV)-induced gene 3 (Ebi3) and IL-12 α chain p35 [33, 34]. IL-35 is a potent immunosuppressive cytokine produced by T_regs_, regulatory B cells (B_regs_) [35], DCs [36], and to a lesser extent, by endothelial cells, smooth muscle cells, and monocytes [37]. The biological effect of IL-35 is poorly understood. However IL-35 is recognised as a typical anti-inflammatory cytokine, and the predominant mechanism of suppression is associated with its ability to suppress T cell proliferation and effector functions [33, 38]. Given the direct immunosuppressive effect of IL-35, many studies have been conducted to evaluate its role in the development of several diseases. IL-35 can suppress several types of chronic inflammatory diseases such as inflammatory bowel disease [26], and decreased the severity of collagen-induced arthritis in animals via enhancement of IL-10 production [39] and suppression of Th17 cells [40]. In an asthma model, intra-tracheal instillation of IL-35 decreased disease severity by diminishing the Th2 cell counts [41] and by reducing the production of IL-17 [42]. In bacterial infections, Shen and associates [35] found that mice without IL-35 expression demonstrated an improved resistance to infection with the intracellular bacterial pathogen *Salmonella typhimurium*. In addition, IL-35 has been increased in the serum of adults and children with sepsis, and administration of anti-IL-35 p35 antibodies diminished dissemination of the bacteria in septic animals [43]. Similarly, tuberculous patients exhibited an increase in serum IL-35 and in mRNA expression of both subunits of IL-35 (p35 and EBI3) in white blood cells and peripheral blood mononuclear cells [44]. However, the role of IL-35 in mycetoma pathogenesis has not been highlighted yet.

With this background, this study was set to determine the IL-1 cytokine family (IL-1β, IL-37) and IL-12 cytokine family (IL-12, IL-35) circulating levels of in patients infected with *M. mycetomatis*, and to explore the association between the cytokine levels and the patients’ demographic characteristics.

## Materials and Methods

### Study population

This case-control study was conducted at the Mycetoma Research Centre, Soba University Hospital, University of Khartoum, Sudan. After a written informed consent, blood samples were taken from patients and a matched control population living in the mycetoma endemic areas of Sudan between 2015 and 2016. Samples collection was previously described in details by Nasr and associates, 2016 [17].

In this study 140 individuals were enrolled; 49 (35%) were females, and 91 (65%) were males with an overall median age of 25 years (range 12–70 years). Seventy patients were infected with *M. mycetomatis*. The study population was divided into two groups; group I: healthy controls [n = 70; median age 25 years (range 12 to 70 years)]. Group II: mycetoma patients [n = 70 patients; median age 25 (range 13 to 70 years)].

The diagnosis of eumycetoma was established by various techniques, and that included imaging, molecular and histopathological techniques, and grain culture [1, 45]. Surgical biopsies are obtained by a wide local incision under anaesthesia and appropriate surgical conditions as part of the routine patients’ treatment protocol [45].

After medical examination, healthy controls were selected from blood bank donors or healthy volunteers to match the patient's birthplace geographically. All healthy controls were questioned for acute or chronic infectious diseases, autoimmune family history and genetic disorders. Then all study participants gave their informed written consent.

### Sample collection

One hundred μl of blood was collected on Whatman qualitative filter paper, Grade 1, circles, diam. 42.5 mm (Sigma-Aldrich Chemical Co., St. Louis, MO, USA) for the determination of cytokines. The use of filter paper dried whole blood spots (DBS) for specimen collection was preferred to facilitate collection, storage and transportation of specimens and it is in line with the World Health Organization recommendations and also used in several previous studies [46–48]. Sera were extracted from filter-paper samples as described previously in details [17].

### IL-1β, IL-37, IL-12 and IL-35 measurement

IL-1β, IL-37, and IL-12 were measured in the sera using commercially available enzyme-linked immunosorbent assay (ELISA) kits (abcam®, Cambridge, UK). Serum levels of IL-35 were estimated using a sandwich ELISA commercial kit (Colorful Gene Biological Technology, Wuhan, China). Cytokine assays were performed in duplicates according to the manufacturers’ protocols. The sensitivity of Human ELISA kits for IL-1β, IL-37, IL-12 and IL-35 cytokines was 0.5 pg/ml.

### Statistical analysis

The data were managed by SPSS version 24.0 statistical software for Windows (IBM© SPSS© statistics) and appropriate statistical tests were used. The results are expressed as mean ± standard deviation (SD) or median with interquartile range (IQR). Spearman correlation test was used to evaluate the associations between serum IL-37 levels and laboratory values as well as serum cytokine levels. For non-parametric data, comparisons between the groups were performed using the Kruskal–Wallis test. One-way ANOVA was used for parametric data. General linear models were used to assess the risk factors for circulating IL-37 pg/ml and IL-35 among mycetoma patients with different disease duration and lesions size of mycetoma infection adjusted with other variables. A test with a *p-*value <0.05 was considered statistically significant.

### Ethical considerations

This study was approved by the Ethics Committee of Soba University Hospital, Khartoum, Sudan. Written informed consent was taken from all the participants before enrolment in the study. The work described here was performed in accordance with the Declaration of Helsinki [49].

## Results

This study included 70 confirmed mycetoma patients and 70 healthy controls; gender and age-matched. Fifty-six patients (80%) were males and 14 (20%) were females. Their ages ranged between 12 and 77 years, and the median age was 25.5 years. The common age group [22 (31.4%)] was 19-24 years, 17 (22.9%) in the age group (25-29) years, and nine (12.9%) were 40 years old or more (Table 1). In this study, a total of 40 subjects [23 patients (32.9%) and 17 (24.3%) healthy controls] were domestic workers, 40 individuals (17 (24.3%) patients and 23 (32.9%) healthy controls] were students and 24 individuals [12 (17.1%) patients and 12 (17.1%) healthy controls] were farmers. Due to the prolonged illness and disability, 12 (17.1%) patients had lost their jobs while only five (7.1%) healthy controls were unemployed. A total of six individuals [5 (7.1%) patients and 1(1.4%) healthy controls] of the patients were housewives. A total of 13 individuals [1(1.4%) patients and 12 (17.1%) healthy controls] were employers (Table 1).

**Table 1.** The Demographic characteristics of the study populations

|  |  | **Controls**<br>**n=70 (%)** | **Mycetoma Patients**<br>**n=70 (%)** |
| --- | --- | --- | --- |
| **Gender** | Female | 14 (20) | 14 (20%) |
|  | Male | 56 (80) | 56 (80) |
| **Age groups**<br>**(years)** | 12 - 18 | 9 (12.9) | 9 (12.9) |
|  | 19 - 24 | 22 (31.4) | 22 (31.4) |
|  | 25 - 29 | 16 (22.9) | 16 (22.9) |
|  | 30 - 39 | 14 (20.0) | 14 (20.0) |
|  | ≥40 | 9 (12.9) | 9 (12.9) |
| **Occupation** | Worker | 17 (24.3) | 23 (32.9) |
|  | Student | 23 (32.9) | 17 (24.3) |
|  | Farmer | 12 (17.1) | 12 (17.1) |
|  | Jobless | 5 (7.1) | 12 (17.1) |
|  | House-wife | 1 (1.4) | 5 (7.1) |
|  | Employer | 12 (17.1) | 1 (1.4) |
| **Discharge\*** | No |  | 35 (50) |
|  | Yes |  | 35 (50) |
| **Lesion**<br>**diameter\*** | Less than 5cm |  | 13 (18.6) |
|  | 5-10cm |  | 22 (31.4) |
|  | More than 10cm |  | 35 (50.0) |
| **Duration\*** | ≤1 year |  | 22 (31.4) |
|  | 2 – 4 years |  | 26 (37.1) |
|  | ≥5 years |  | 22 (31.4) |
| **Medication\*** | Itraconazole<br>200mg/day |  | 46 (65.7) |
|  | Ketoconazole<br>400mg/day |  | 24 (34.3) |

\*These parameters only for mycetoma patients.

### Correlations between cytokine levels (IL-1β, IL-37, IL-12 and IL-35) stratified by the lesion diameter among mycetoma patients and control group

The levels of cytokines (IL-1β, IL-37, IL-12 and IL-35) were constitutively correlated among mycetoma patients with different lesions diameters. The levels of IL-1β were constitutively positively correlated with IL-12 and lesions diameter, (Table 2). On the other hand, the levels of cytokine IL-1β were constitutively negatively correlated with IL-37 and IL-35, (Table 2). Furthermore, the levels of cytokine IL-37 were constitutively positively correlated with IL-35 (Table 2). However, the levels of cytokine IL-37 and IL-35 were constitutively negatively correlated with IL-12 (Table 2).

**Table 2.** Correlations between the serum levels of (IL-1β, IL-37, IL-12 and IL-35) and the lesion diameter and the control group

| Lesion diameter | Cytokine Levels pg/ml | IL-1β pg/ml | IL-37 pg/ml | IL-12 pg/ml | IL-35 pg/ml |
| --- | --- | --- | --- | --- | --- |
| **Controls (no lesion)** | IL-1β | 1 | | | |
|  | IL-37 | 0.115 | 1 |  |  |
|  | IL-12 | 0.137 | -0.023 | 1 |  |
|  | IL-35 | -0.125 | 0.037 | 0.065 | 1 |
| **$\leq 5$cm** | IL-1β | 1 | | | |
|  | IL-37 | -0.992** | 1 |  |  |
|  | IL-12 | 0.987** | -0.996** | 1 |  |
|  | IL-35 | -0.924** | 0.945** | -0.949** | 1 |
| **5-10cm** | IL-1β | 1 | | | |
|  | IL-37 | -0.991** | 1 |  |  |
|  | IL-12 | 0.991** | -0.995** | 1 |  |
|  | IL-35 | -0.983** | 0.989** | -0.983** | 1 |
| **$\geq 10$cm** | IL-1β | 1 | | | |
|  | IL-37 | -0.998** | 1 |  |  |
|  | IL-12 | 0.969** | -0.971** | 1 |  |
|  | IL-35 | -0.994** | 0.995** | -0.964** | 1 |
| ** Spearman's rho Correlation is significant at the 0.01 level (2-tailed). |  |  |  |  |  |

### Correlations between cytokine levels (IL-1β, IL-37, IL-12 and IL-35) stratified by the disease duration and control group

In the patients' group, the levels of cytokines (IL-1β, IL-37, IL-12 and IL-35) were constitutively correlated with the duration of mycetoma infection. Levels of IL-1β showed a consistent positive correlation with IL-12 and negative correlation with IL-37 and IL-35, (Table 3). Whereas, levels of IL-37 were constitutively positively correlated with IL-35 (Table 3). However, the levels of IL-37 and IL-35 were constitutively negatively correlated with IL-12, (Table 3).

**Table 3.** Correlations between the serum levels of (IL-1β, IL-37, IL-12 and IL-35) and the duration of mycetoma infection and the control group

| Duration | Cytokine Levels<br>pg/ml | IL-1β pg/ml | IL-37 pg/ml | IL-12 pg/ml | IL-35 pg/ml |
| --- | --- | --- | --- | --- | --- |
| **Control** | IL-1β | 1 | | | |
|  | IL-37 | 0.115 | 1 |  |  |
|  | IL-12 | 0.137 | -0.023 | 1 |  |
|  | IL-35 | -0.125 | 0.037 | 0.065 | 1 |
| **≤1 Years** | IL-1β | 1 | | | |
|  | IL-37 | -0.997** | 1 |  |  |
|  | IL-12 | 0.997** | -0.999** | 1 |  |
|  | IL-35 | -0.996** | 0.997** | -0.995** | 1 |
| **2 – 4 years** | IL-1β | 1 | | | |
|  | IL-37 | -0.999** | 1 |  |  |
|  | IL-12 | 0.996** | -0.997** | 1 |  |
|  | IL-35 | -0.997** | 0.997** | -0.994** | 1 |
| **≥5 years** | IL-1β | 1 | | | |
|  | IL-37 | -0.999** | 1 |  |  |
|  | IL-12 | 0.986** | -0.986** | 1 |  |
|  | IL-35 | -0.998** | 0.997** | -0.986** | 1 |
| ** Spearman's rho Correlation is significant at the 0.01 level (2-tailed). |  |  |  |  |  |

### Analysis of the serum levels of IL-1β, IL-37, IL-12 and IL-35 within the different lesion diameters among mycetoma patients and control groups

Circulating serum cytokine levels were determined in all mycetoma patients and were compared between the different lesion diameters among mycetoma patients and healthy controls. Overall, there was a significant difference in the levels of all studied cytokines between the four groups (Three levels of lesion diameter and the healthy controls), (Table 4). Distribution of IL-1β levels has decreased dramatically with lesion diameter [for lesion diameter ≤ 5 cm: the mean ± SD (3.39 ± 1.07); for 5-10 cm: (2.32 ± 0.05); for ≥ 10 cm: (2.08 ± 0.11), *p*-value < 0.001)]. However, the circulating serum levels of IL-37 were significantly increased with lesion diameter [for lesion diameter ≤ 5 cm: the mean ± SD (107.92 ± 5.96); for 5-10 cm: (141.45 ± 12.96) and for ≥ 10 cm: (193.20 ± 15.01), *p*-value < 0.001)] (Table 4). Our results showed a significant reduction of circulating IL-12 levels versus lesion diameter [for lesion diameter ≤ 5 cm: the mean ± SD (25.22±3.34); for 5-10 cm: (14.45± 3.32); for ≥ 10 cm: (9.65 ± 0.36), *p* value < 0.001)], (Table 4). Circulating levels of IL-35 were significantly increased with increasing lesions’ diameter [for lesion diameter ≤ 5 cm: the mean ± SD (255.15 ± 1.72); for 5-10 cm: (263.23 ± 3.26); ≥ 10 cm: (449.71 ± 22.2), (*p* value < 0.001)], (Table 4).

**Table 4.** Analysis of the serum cytokine levels of (IL-1β, IL-37, IL-12 and IL-35) within the different lesion diameter among mycetoma patient and control groups

| Cytokines Levels pg/ml | Lesion diameter | Mean± SD | Median | (Q1-Q3) | P-value* |
| --- | --- | --- | --- | --- | --- |
| **IL-1β** | **Control** | 1.16± 3.15 | 0.11 | 0.3- 0.6 | **<0.001** |
| | **≤5cm** | 3.39± 1.07 | 2.45 | 2.5-4.5 | |
| | **5-10cm** | 2.32± 0.05 | 2.3 | 2.3- 2.3 | |
| | **≥10cm** | 2.08± 0.11 | 1.97 | 2.1-2.1 | |
| **IL-37** | **Control** | 22.06± 2.39 | 20 | 22-24 | **<0.001** |
| | **≤5cm** | 107.92± 5.96 | 104 | 108-112 | |
| | **5-10cm** | 141.45± 12.96 | 129 | 143-152 | |
| | **≥10cm** | 193.20± 15.01 | 184.17 | 189-199 | |
| **IL-12** | **Control** | 2.46± 1.02 | 1.96 | 2.4-2.5 | **<0.001** |
| | **≤5cm** | 25.22± 3.34 | 23.2 | 25-28 | |
| | **5-10cm** | 14.45± 3.32 | 12.4 | 12.9- 18.2 | |
| | **≥10cm** | 9.65± 0.36 | 9.49 | 9.5- 9.8 | |
| **IL-35** | **Control** | 15.97± 2.6 | 14.6 | 16.4-18.2 | **<0.001** |
| | **≤5cm** | 255.15± 1.72 | 253 | 255-257 | |
| | **5-10cm** | 263.23± 3.26 | 261 | 262-267 | |
| | **≥10cm** | 449.71± 22.2 | 430 | 447-456 | |

\*P values are derived from non-parametric method; *Kruskal Wallis* test.

### Analysis of the serum levels of IL-1β, IL-37, IL-12 and IL-35 among mycetoma patients stratified by different durations of mycetoma infection compared to the control group

Circulating levels of IL-1β had significantly decreased with increasing disease duration [(≤ 1 year; median = 2.3 pg/ml), (2-4 years; median = 2.2 pg/ml) and (≥ 5 years; median = 2.2 pg/ml)], *p v*alue = 0.017 (Table 5). Serum levels of IL-12 dramatically decreased with the increase in disease duration [(≥ 1 year; median = 12.5 pg/ml), (2-4 years; median = 10.2 pg/ml) and (≥ 5 years; median = 9.8 pg/ml)] and *p* value < 0.001), (Table 5). However, circulating levels of IL-37 were positively increased with different durations of mycetoma infection [(≤ 1 year; median = 145 pg/ml), (2-4 years; median = 178 pg/ml) and (≥ 5 years; median = 185.2 pg/ml)], p value <0.001, (Table 5). Similarly, serum levels of IL-35 were also significantly increased with increasing duration of mycetoma infection [(≤ 1 year; median = 262.5 pg/ml), (2-4 years; median = 423.5 pg/ml) and (≥ 5 years; median = 436 pg/ml)], *p* value < 0.001, (Table 5).

**Table 5.** Analysis of the serum cytokine levels of (IL-1β, IL-37, IL-12 and IL-35) among mycetoma patient stratified by different duration of mycetoma infection compared to controls group

| Cytokines<br>pg/ml | Duration | Mean± SD | Median | (Q1- Q3) | P-value* |
| --- | --- | --- | --- | --- | --- |
| **IL-1β** | Control | 1.2± 3.1 | 0.4 | (0.1- 0.6) | 0.017 |
| | ≤1 year | 2.3± 0.1 | 2.3 | (2.2- 2.4) | |
| | 2 - 4 years | 2.6± 0.9 | 2.2 | (2.1- 2.4) | |
| | ≥5 years | 2.3± 0.5 | 2.2 | (2.0- 2.3) | |
| **IL-37** | Control | 22.1± 2.4 | 22.0 | (20- 24) | <0.001 |
| | ≤1 year | 146.7± 28.4 | 145.0 | (122- 160) | |
| | 2 - 4 years | 160.9± 41.3 | 178.0 | (127- 190) | |
| | ≥5 years | 175.7± 33.9 | 185.2 | (156- 194) | |
| **IL-12** | Control | 2.5± 1.0 | 2.4 | (2.0- 2.5) | <0.001 |
| | ≤1 year | 14.9± 5.0 | 12.5 | (10.5- 19.5) | |
| | 2 - 4 years | 15.0± 7.5 | 10.2 | (9.5- 19.4) | |
| | ≥5 years | 12.1± 5.4 | 9.8 | (9.5- 12.1) | |
| **IL-35** | Control | 16.0± 2.6 | 16.4 | (14.6- 18.2) | <0.001 |
| | ≤1 year | 301.9± 75.9 | 262.5 | (260- 271) | |
| | 2 - 4 years | 363.8± 101.5 | 423.5 | (260- 447) | |
| | ≥5 years | 397.6± 88.1 | 436.0 | (267- 450) | |

\*P values are derived from non-parametric method; *Kruskal Wallis* test.

### Risk factors for increased IL-37 levels in mycetoma patients with different lesion diameters

The analysis of the risk factors of higher levels of IL-37 in patients of mycetoma showed a significant negative association with IL-1β, where a unit increment in IL-1β decreases the levels of IL-37 by 9.1 pg/ml, *p*-value = 0.008, (Table 6). Serum levels of IL-37 among the patients with lesion diameter ≤ 5 cm and 5-10 cm have significantly lower on average by 75.4% and 52.6%, respectively, compared to patients with lesion diameter ≥ 10 cm (reference group). Serum levels of IL-37 among the patients of mycetoma showed no significant difference between males and females, *p-value* = 0.176, (Table 6). Circulating levels of IL-37 significantly decreased with increasing age groups [(19-24 years); *p-*value = 0.010 and (30-39) years; p-value = 0.029)] (Table 6).

**Table 6.** Risk factors for circulating cytokines IL-37 pg/ml in mycetoma patients with different lesion diameters

| variable | Category | B<sup>‡</sup> | 95% Confidence Interval |  | p-value |  |
| --- | --- | --- | --- | --- | --- | --- |
|  |  |  | (Lower to Upper) |  |  |  |
| Intercept |  | 228.4 | (208.6 to 248.1) |  | <0.001 |  |
| IL-1β |  | -9.1 | (-15.7 to -2.5) |  | 0.008 |  |
| Lesion diameter | ≤5cm | -75.4 | (-88.4 to -62.3) |  | <0.001 |  |
|  | 5-10cm | -52.6 | (-64.3 to -40.8) |  | <0.002 |  |
|  | ≥10cm | 0 |  |  |  |  |
| Gender | Female | -5.3 | (-13.1 to 2.5) |  | 0.176 |  |
|  | Male | 0 |  |  |  |  |
| Age groups years | 12 - 18 | -11.7 | (-23.2 to -0.2) |  | 0.047 |  |
|  | 19 - 24 | -13.3 | (-23.2 to -3.3) |  | **0.010** |  |
|  | 25 - 29 | -9.5 | (-20.0 to 1.1) |  | 0.079 |  |
|  | 30 - 39 | -11.8 | (-22.3 to -1.2) |  | **0.029** |  |
|  | ≥40 | 0 |  |  |  |  |
| Medication | Itraconazole 200mg/day | -5.7 | (-16.4 to 5) |  |  | 0.293 |
|  | Ketoconazole 400mg/day | 0 |  |  |  |  |

<sup>‡</sup>B (95%CI) adjusted with lesion diameter, gender, age groups and medical treatments.

### Risk factors for increasing circulating IL-35 and the patients’ demographic characteristics and lesion diameters

The analysis of the risk factors of higher serum levels of IL-35 in mycetoma patients showed no significant association with IL-12, *p*-value = 0.182, (Table 7). Circulating levels of IL-35 among the patients with lesions’ diameter ≤ 5 cm and 5-10 cm were significantly decreased, *p*-value < 0.001) by 174.4% and 176.5, respectively, compared to patients with lesion diameter ≥ 10 cm (reference group) (Table 7).

**Table 7.**
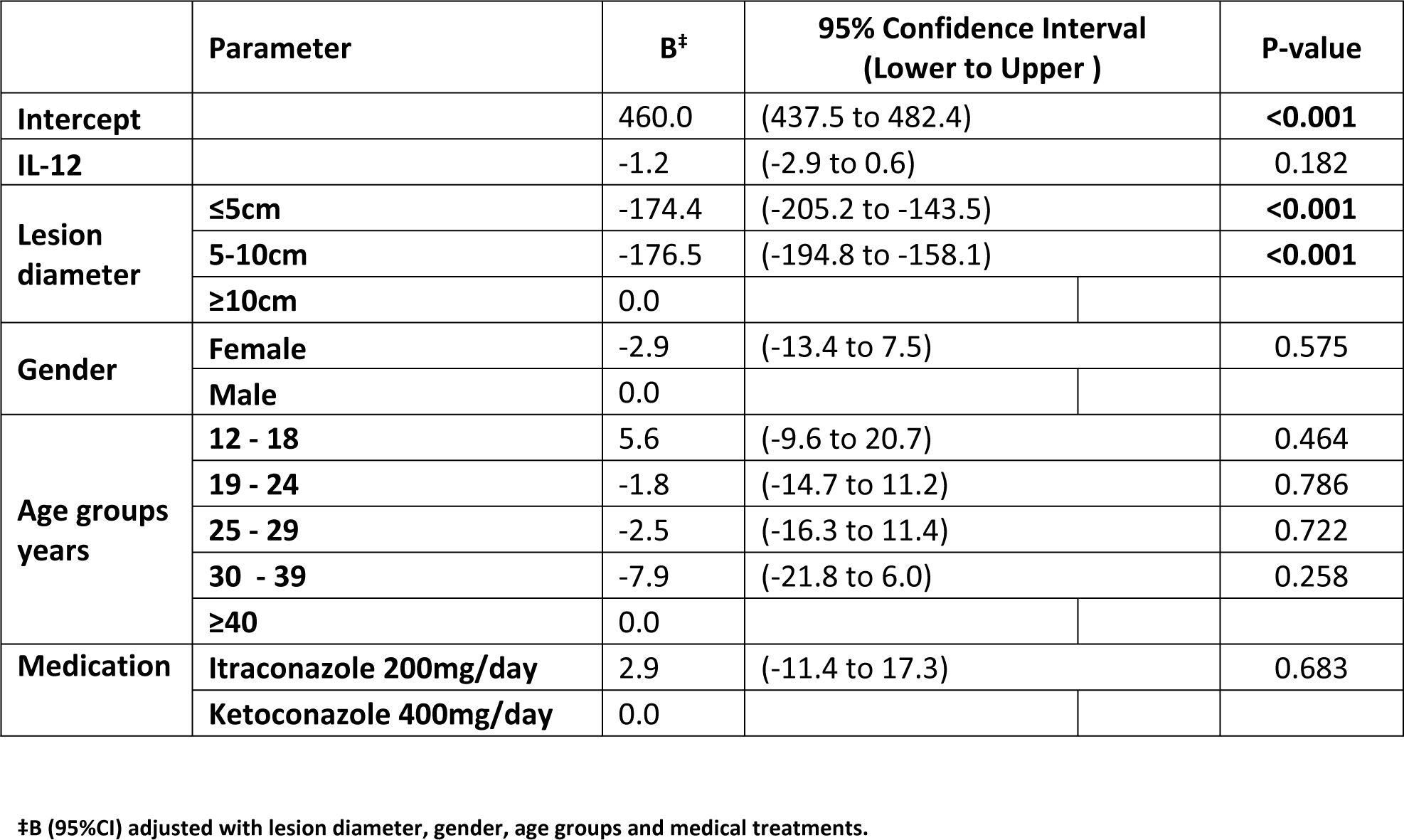
Risk factors for circulating cytokines IL-35 pg/ml in mycetoma patients with different lesion diameters

Serum levels of IL-35 among mycetoma patients showed no significant difference between males and female, *p*-value = 0.575, (Table 7). Circulating levels of IL-35 showed no significant association with the different age groups and different types of antifungal medication given (Table 7).

### Risk factors for circulating IL-37 and the patients’ demographic characteristics and disease duration

The analysis of the risk factors of higher circulatory levels of IL-37 in patients of mycetoma showed a significant negative association with IL-1β, where a unit increment in IL-1β decreases the levels of IL-37 by 35.28 pg/ml, *p* < 0.001 (Table 8). Levels of IL-37 among patients with a disease duration ≤ one year had significantly decreased on average by 18.45 compared to patients with a disease duration ≥ 5years (reference group). However, there was no significant difference in levels of IL-37 between patients with infection duration 2-4 years and ≥ five years, *p*-value = 0.793. Also, serum levels of IL-37 among mycetoma patients showed no significant difference between males and females, *p*-value = 0.627 (Table 8). Furthermore, the circulating levels of IL-37 had significantly decreased with increasing age groups [(19-24 years; *p*-value = 0.0100), 25-29 years; *p*-value =0.030 and (30- 39) years; *p*-value = 0.022)] (Table 8). Interestingly, levels of IL-37 among mycetoma patients showed a statistically significant difference between Itraconazole compared to Ketoconazole, p < 0.001, (Table 8).

**Table 8.** Risk factors for circulating cytokines IL-37 pg/ml among mycetoma patients with different duration of mycetoma infection

|  | Parameter | B<sup>‡</sup> | 95% Confidence Interval |  | P-value |
| --- | --- | --- | --- | --- | --- |
|  |  |  | Lower to Upper |  |  |
| Intercept |  | 254.499 | 225.3 to 283.7 |  | <0.001 |
| IL-1β |  | -35.28 | -43.9 to -26.7 |  | <0.001 |
| Duration of mycetoma infection | ≤1 year | -18.45 | -32.3 to -4.6 |  | 0.010 |
|  | 2 - 4 years | -1.774 | -15.2 to 11.7 |  | 0.793 |
|  | ≥5 years | 0 |  |  |  |
| Gender | Female | -3.223 | -16.4 to 10.0 |  | 0.627 |
|  | Male | 0 |  |  |  |
| Age groups years | 12 - 18 | -14.631 | -35.6 to 6.4 |  | 0.169 |
|  | 19 - 24 | -20.445 | -37.8 to -3.1 |  | 0.022 |
|  | 25 - 29 | -20.199 | -38.5 to -2.0 |  | 0.030 |
|  | 30 - 39 | -22.68 | -42.0 to -3.4 |  | 0.022 |
|  | ≥40 | 0 |  |  |  |
| Medication | Itraconazole 200mg/day | 23.977 | 11.8 to 36.1 |  | <0.001 |
|  | Ketoconazole 400mg/day | 0 |  |  |  |

<sup>‡</sup>B (95%CI) adjusted with Duration of mycetoma infection, gender, age groups and medical treatments.

### Risk factors for circulating IL-35 among mycetoma patients

The analysis of the risk factors of higher levels of IL-35 in patients of mycetoma revealed a significant negative association with IL-12; as a unit increment in IL-12 decreases the levels of IL-35 by 8.99 pg/ml, *p*-value < 0.001 (Table 9). Levels of IL-35 among the patients with a mycetoma with a disease duration of ≤ 1 year had significantly decreased, *p* value= 0.002), on average by 41.82 pg/ml compared to patients with a disease duration ≥ 5years (reference group) (Table 9). However, patients with an infection duration of 2-4 years and ≥ five years showed no significant difference in IL-35 levels, *p*-value = 0.391. Furthermore, there was no significant difference (*p*-value = 0.49) in IL-35 levels between male and female mycetoma sufferers (Table 9). Additionally, the circulating levels of IL-35 showed no significant association with the different age groups (Table 9). Interestingly, treatment with Itraconazole significantly increased circulating levels of IL-35 among mycetoma patients compared to treatment with Ketoconazole, *p*-value <0.001, (Table 9).

**Table 9.** Risk factors for circulating cytokines IL-35 pg/ml among mycetoma patients with different duration of mycetoma infection

|  | Parameter | B<sup>‡</sup> | 95% Confidence Interval |  | P-value |
| --- | --- | --- | --- | --- | --- |
|  |  |  | (Lower to Upper ) |  |  |
| Intercept |  | 436.50 | 394.36 to 478.64 |  | <0.001 |
| IL-12 |  | -8.99 | -10.73 to -7.25 |  | <0.001 |
| Duration of mycetoma infection | ≤1 year | -41.82 | -67.31 to -16.32 |  | 0.002 |
|  | 2 - 4 years | -10.64 | -35.28 to 14.01 |  | 0.391 |
|  | ≥5 years | 0 |  |  |  |
| Gender | Female | -8.47 | -32.85 to 15.90 |  | 0.49 |
|  | Male | 0 |  |  |  |
| Age groups years | 12 - 18 | 9.23 | -29.02 to 47.48 |  | 0.631 |
|  | 19 - 24 | 8.00 | -24.05 to 40.06 |  | 0.619 |
|  | 25 - 29 | -2.19 | -36.43 to 32.05 |  | 0.899 |
|  | 30 - 39 | -30.58 | -66.01 to 4.84 |  | 0.089 |
|  | ≥40 | 0 |  |  |  |
| Medication | Itraconazole 200mg/day | 101.17 | 78.40 to 123.95 |  | <0.001 |
|  | Ketoconazole 400mg/day | 0 |  |  |  |

<sup>‡</sup>B (95%CI) adjusted with Duration of mycetoma infection, gender, age groups and medical treatments.

## Discussion

Although mycetoma represents a major health problem in many tropical and subtropical areas, there are no prevention or control measures for this neglected disease [22, 23]. In mycetoma endemic areas, most individuals have antibodies against the causative agents. However only a few develop the disease [4]. Few researchers believed that patients who develop mycetoma seem to be deficient in their cell-mediated immunity [6]. Hence, we aimed to investigate the profiles of the pro-inflammatory (IL-1β and IL-12) and the anti-inflammatory immunosuppressive (IL-37 and IL-35) cytokines among mycetoma patients and their association with disease characteristics. As far as we know, this is the first study addressing the relation between immunosuppressive cytokines and mycetoma infection. Our data showed that eumycetoma patients presented higher circulating levels of IL-1β, IL-12, IL-37 and IL-35 compared to controls. Moreover, serum levels of IL-1β and IL-12 were significantly decreased with increasing lesions’ diameter and disease duration, whereas levels of IL-37 and IL-35 were significantly higher with increasing lesions’ diameter and disease duration. These findings indicate that immunosuppressive cytokines like IL-37 and IL-35, which could suppress cell-mediated immune responses, may have a negative impact on the disease progression.

Data of the current study clearly showed the serum levels of IL-1β and IL-12 in eumycetoma patients with different lesion size and disease duration were positively correlated with each other, and negatively correlated with IL-37. It has been demonstrated that the first line of the innate immune response against mycetoma infection is by phagocytes, from which macrophages represent the major phagocytic cells [50]. In general, protective immunity to fungal infections [51] involves activation of Toll-like receptors (TLRs) generating inflammatory cytokines through pattern-recognition receptors and pathogen-associated molecular patterns [52, 53]. IL-1β and other pro-inflammatory cytokines are produced early in response to fungal infections and promote phagocytosis and other means of the innate immune response [54, 55]. Following inflammatory stimuli, several cell types including immune and non-immune cells produce IL-37, as a protective mechanism to prevent runaway inflammation and excessive tissue damage [56]. IL-37 directly inhibits generation of pro-inflammatory cytokines and down-regulates macrophage cytokine release, and therefore innate immunity [57, 58]. Moreover, IL-37 induces macrophages towards an M2-like phenotype [59]. M1 macrophages are the most critical effector cells in the innate immune defence system and are characterised by high expression levels of iNOS, subsequent NO production and secretion of pro-inflammatory cytokines, such as IL-1β and IL-12 [60]. However, M2 macrophages secrete anti-inflammatory cytokines, such as IL-10 [61] and express arginase 1, which inhibits NO production, thus rendering these cells ineffective in killing infectious agents including fungal agents [61, 62]. Furthermore, DCs expressing IL-37 secreted higher levels of IL-10 and reduced levels of IL-1β and IL-12. Therefore, the presence of IL-37 in DCs impairs their function in prime T cells and promotes their ability to induce Treg cells that produce IL-10, which is also a potent anti-inflammatory cytokine [30].

Our results have consistently shown higher circulatory levels of IL-37 in patients of mycetoma which is negatively associated with IL-1β, as a unit increment in IL-1β decreases the levels of IL-37 by 35.28 pg/ml. Based on the aforementioned data, we can speculate that IL-37 could play a role in damping inflammatory response in mycetoma infection which leads to disease progression and this is not in the patient’s favour.

In the current work, the circulating levels of IL-1β and IL-12 in eumycetoma patients with different lesion size and disease duration were negatively correlated with IL-35; whereas serum levels of IL-35 were increased with increasing lesion size and disease duration, and levels of IL-1β and IL-12 simultaneously decreased. This may probably be an attempt to dampen ongoing inflammation. Both IL-1β and IL-12 have a pivotal role in inflammatory and cell-mediated immune responses. Macrophages, Th1 and cytotoxic T-cells (CTLs), which constitute the main component of cell-mediated immunity, play an important role in the protective immunity against mycetoma infection [2]. As fatal dissemination of *N. asteroides* infection occurs in nude athymic rats and mice [11, 12], T cells from previously immunised animals are able to kill *N. asteroides* in new infections [12, 13]. IL-35 could suppress Th1 and macrophage responses [63], whereas deficiency in IL-35 increases macrophage’s activation and induces Th1 responses [35, 63]. The increased immunity found in mice lacking IL-35 is associated with higher activation of macrophages and inflammatory T cells, as well as enhancing the function of antigen-presenting cells [35]. In another infection model, Cao and his co-workers reported higher serum levels of IL-35 in septic patients compared to controls, and IL-35 gradually increased with increased sepsis severity.

Moreover, administration of anti-IL-35 antibodies diminished dissemination of the bacteria in septic animals and enhanced local neutrophil recruitment with increases in inflammatory cytokines and chemokines production [43]. Furthermore, IL-35 suppressed the proliferation of antigen-specific CTLs and IFN-γ production [64]. Our data revealed that higher levels of IL-35 in patients with mycetoma is negatively associated with IL-12, where a unit increment in IL-12 decreases the levels of IL-35 by 8.99 pg/ml. This finding indicates that IL-35 may be a risk factor for mycetoma infection and have a negative role in the clinical presentation of the disease.

Prevalence of mycetoma infection may vary with age. Data from this study showed a variation of mycetoma prevalence with age; 74.3 % of the patients’ age was 19-39 years. This finding is consistent with previous studies which reported that mycetoma mostly affects ages between 20 and 40 years. Our data also demonstrated that mycetoma infection was predominant in males, as the male to female ratio in patient’s group was 4:1. This finding is running parallel with the results of a previous study which demonstrated that in a tertiary facility in Khartoum, Sudan, the male to female ratio is 4:1, whereas at the primary care level in White Nile State, Sudan, the reported male to female ratio was 1.6:1. Another studies reported that male to female ratios in mycetoma infections was in the range of 1.6-6.6:1 [2]. The predominance of mycetoma in males may be attributed to increased exposure in men who engage in different manual labours including agricultural work. Moreover, the influence of sex hormones might have a role in susceptibility to mycetoma infections and disease progression [4, 65].

Our data showed that about, 50 % of the patients have lesion diameter more than 10 cm. This result reflected that most mycetoma patients tend to present late with massive lesions. This finding could be attributed to the nature of mycetoma which is usually painless and slowly progressive. In addition, the lack of health facilities in endemic areas, the low socio-economic status of the affected patients and their poor health education [1, 4, 66] are amongst the reasons why the current treatment of mycetoma is suboptimal, characterised by low cure rates and frequent recurrence often leading to amputation [67, 68]. However, clinical experience shows that early and small mycetoma lesions are associated with good outcome and prevent severe complications of the disease.

One of the remarkable findings of the current study is the significant increase of IL-37 and IL-35 levels with Itraconazole treatment compared to the Ketoconazole. A previous study by Friccius and colleagues suggested that the dose of 10 μg/ml Itraconazole leads to strong inhibition of the cytokines IL-2, IL-4, IL-9 and IFN-γ and slight inhibition of TNF-α cytokine production in PBMC after 6 and 24 hours of incubation. These results demonstrate that IL-35 and IL-37 can be one of the underline factors associated with inhibition of the cytokines related to Itraconazole [69].

In conclusion, our study revealed that the levels of IL-37 and IL-35 were consistently positively correlated with different diameters of mycetoma lesions as well as its duration. However, the levels of IL-1β and IL-12 were consistently negatively correlated with different diameters of lesions and the duration of mycetoma infection. The analysis of the risk factors of higher circulatory levels of IL-37 in patients of mycetoma showed a negative significant association with IL-1β cytokine, where a unit increment in IL-1β will decrease the levels of IL-37 by 35.28 pg/ml. Levels of IL-37 among the patients with a mycetoma infection duration ≤ one year had significantly decreased on average by 18.45 compared to patients with a mycetoma infection duration ≥ 5years (reference group). Furthermore, the risk factors of higher levels of IL-35 in patients of mycetoma revealed a significant negative association with IL-12, as a unit increment in IL-12 decreases the levels of IL-35 decrease by 8.99 pg/ml *p* < 0.001. Levels of IL-35 among the patients with a mycetoma infection duration ≤ 1 year had significantly decreased *p*-value = 0.002) on average by 41.82 compared to patients with a mycetoma infection duration ≥ 5years (reference group). More investigations are needed to explore the mechanism by which IL-35 and IL-37 contribute in the mycetoma infection outcomes. This will help in understanding the role of these cytokines IL-35 and IL-37 in the pathogenesis of mycetoma, and may exploit it as a potential therapeutic target to prevent mycetoma diseases recurrence.

## Acknowledgements

The authors are grateful to the patients, controls and their families and the staff at the Mycetoma Research Centre at Soba University Hospital, University of Khartoum-Sudan, for their continued collaboration and generous hospitality during a decade of samples collection. The authors are thankful to Dr Hamza A., Dundee University, the UK for improving the English language and proofreading the Manuscript.

## Supporting Information Legends

STROBE checklist

